# Asymmetric Division Promotes Therapeutic Resistance in Glioblastoma Stem Cells

**DOI:** 10.1101/569962

**Authors:** Masahiro Hitomi, Anastasia P. Chumakova, Daniel J. Silver, Arnon M. Knudsen, W. Dean Pontius, Stephanie Murphy, Neha Anand, Bjarne W. Kristensen, Justin D. Lathia

## Abstract

Asymmetric cell division (ACD) enables the maintenance of a stem cell population while simultaneously generating differentiated progeny. Cancer stem cells (CSCs) undergo multiple modes of cell division during tumor expansion and in response to therapy, yet the functional consequences of these division modes remain to be determined. Using a fluorescent reporter for cell surface receptor distribution during mitosis, we found that ACD in glioblastoma CSCs generated a daughter cell with enhanced therapeutic resistance and increased co-inheritance of epidermal growth factor receptor (EGFR) and neurotrophin receptor (p75NTR). Stimulation of both receptors maintained self-renewal under differentiation conditions. While p75NTR knockdown did not compromise CSC maintenance, therapeutic efficacy of EGFR inhibition was enhanced, indicating that co-inheritance of p75NTR and EGFR promotes resistance to EGFR inhibition through a redundant mechanism. These data demonstrate that ACD produces progeny with co-enriched growth factor receptors, which contributes to the generation of a more therapeutically resistant CSC population.

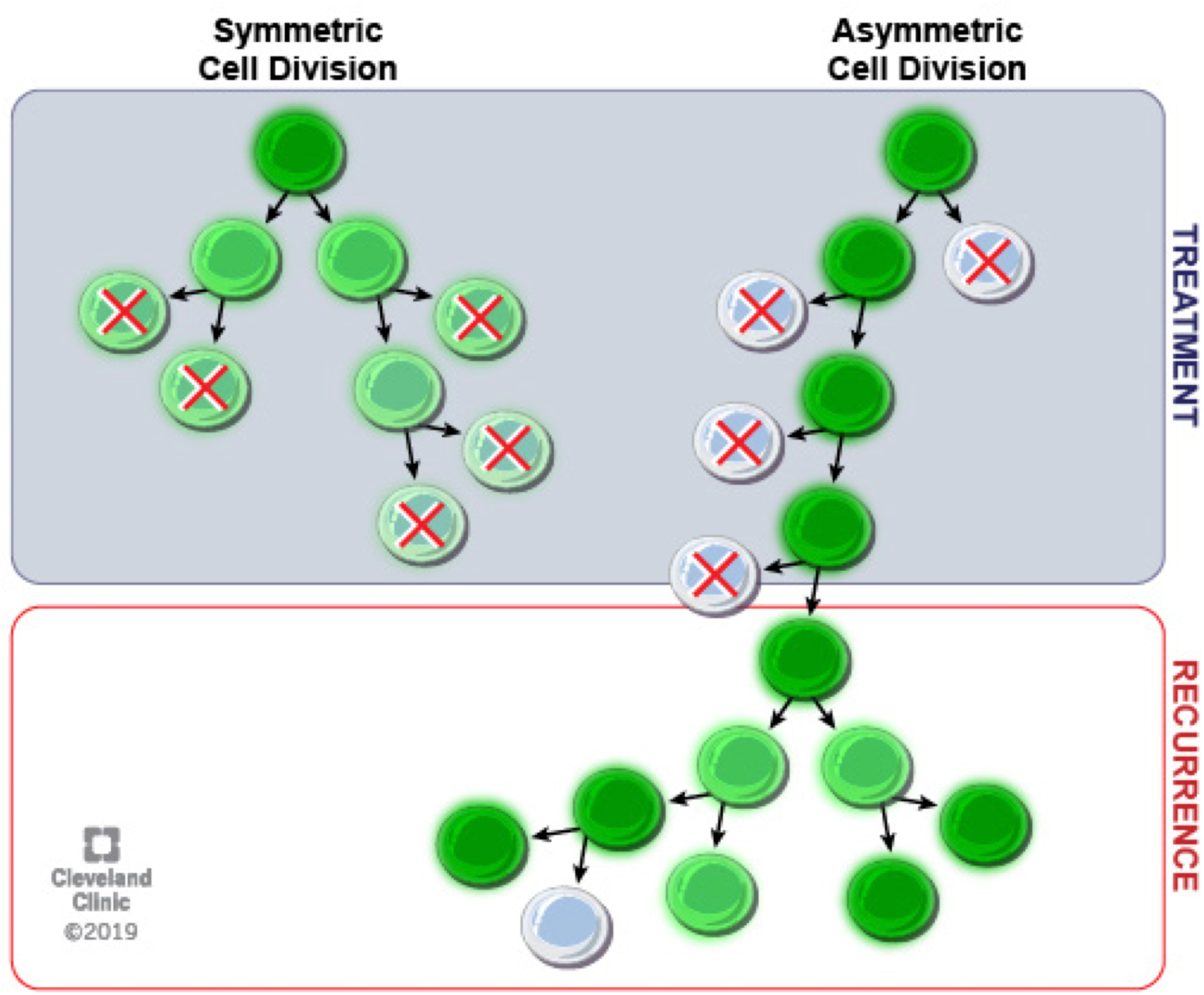
Graphical abstract.

## Introduction

Experimental studies have provided evidence that cancer stem cells (CSC) drive tumor growth and are resistant to conventional therapies (Batlle and Clevers, 2017; Clarke et al., 2006). Therapeutic resistance in CSCs has been attributed to multiple mechanisms, including active drug efflux pumps, enhanced DNA repair capacity, slow proliferation rate, and activation of key survival pathways (Bao et al., 2006; Chen et al., 2016; Moitra et al., 2011). While these mechanisms have been identified, the mechanisms by which CSCs emerge, are maintained, and evolve as a result of therapies have yet to be determined. Central to the identity of CSCs is the ability to execute multiple modes of cell division. Asymmetric cell division (ACD) is a cellular mechanism to generate heterogeneity while simultaneously maintaining a stem cell population(Venkei and Yamashita, 2018). ACD has been observed in multiple advanced cancers (Bu et al., 2016; Lathia et al., 2011; Srinivasan et al., 2016; Wang et al., 2016), yet its functional contribution to tumorigenesis is not well understood. As ACD can enrich fate-determining molecules in one daughter cell, we hypothesized that this cellular mechanism may be leveraged in CSCs to generate therapeutically resistant progeny by concentrating pro-survival molecules to one daughter cell at the expense of the other. We previously demonstrated that CSCs from glioblastoma (GBM), the most common primary malignant brain tumor (Thakkar et al., 2014), execute ACD (Lathia et al., 2011). We now show that a functional consequence of ACD is the ability to enrich pro-survival signaling activity by epidermal growth factor receptor (EGFR) and nerve growth factor receptor (p75NTR) in one daughter cell. Interfering with these signaling mechanisms resulted in the ability to sensitize CSCs to previously ineffective treatment regimens targeting EGFR (Clarke et al., 2014; van den Bent et al., 2009).

## Results and Discussion

### A lipid-raft reporter informs the mode of cell division and fate of the progeny

We previously demonstrated the asymmetric inheritance of CD133, a CSC marker, in a minor fraction of GBM CSC mitoses (Lathia et al., 2011). The frequency of this type of cell division increased under a differentiation-inducing condition, deprivation of growth factors, which also increased the incidence of asymmetric cell fate choice determined by lineage tracing (Lathia et al., 2011). To establish a direct connection between ACD and differential cell fate determination, we developed a green fluorescence protein (GFP)-based reporter for CD133 inheritance at the time of mitosis. Based on the observation that CD133 is enriched in cholesterol-rich lipid rafts (Roper et al., 2000), we reasoned that a fusion protein containing the N-terminus of Lyn fused to GFP (PMGFP) that is enriched in lipid rafts through myristoylation/palmitoylation of its N-terminus (Pyenta et al., 2001) would report CD133 distribution in the two daughter cells during mitosis. To validate the ability of this reporter to mark modes of cell division in dividing CSCs, we stained PMGFP-expressing CSCs for lipid rafts using cholera toxin B (CTB) and for CD133 using immunofluorescence and observed that this reporter co-segregated with both lipid rafts and CD133 during ACD (**Fig. 1A**). Quantification of the fluorescent signals of dividing daughter cells demonstrated a correlation between PMGFP and CD133 asymmetry (**Fig. 1B**), with the majority of cells co-segregating both markers to the same daughter cell. Importantly, the expression of this reporter gene did not adversely impact the self-renewal capacity of CSCs (**Supplemental Fig. 1A**). When combined with time-lapse microscopy and quantitative image analysis at single-cell resolution, this PMGFP reporter system for cell division mode enabled real-time monitoring of the mode of cell division and prospective determination of the fate of the resulting progeny. We measured PMGFP asymmetry during mitosis and traced the daughter cells through the recorded time-lapse images. After the recording, the cells were fixed and stained to assess SOX2 expression as a surrogate for the CSC state (**Fig. 1C**). This approach revealed that the daughter cell expressing higher PMGFP at the time of mitosis also eventually expressed elevated SOX2 compared to its counterpart under a differentiation-inducing condition (**Fig. 1C, Supplemental Fig. 1B**). These results suggest that PMGFP provides a reliable reporting of the asymmetric inheritance of lipid rafts. This in turn predicts the daughter cells favored to maintain the CSC phenotype with high fidelity.

**Figure 1.**
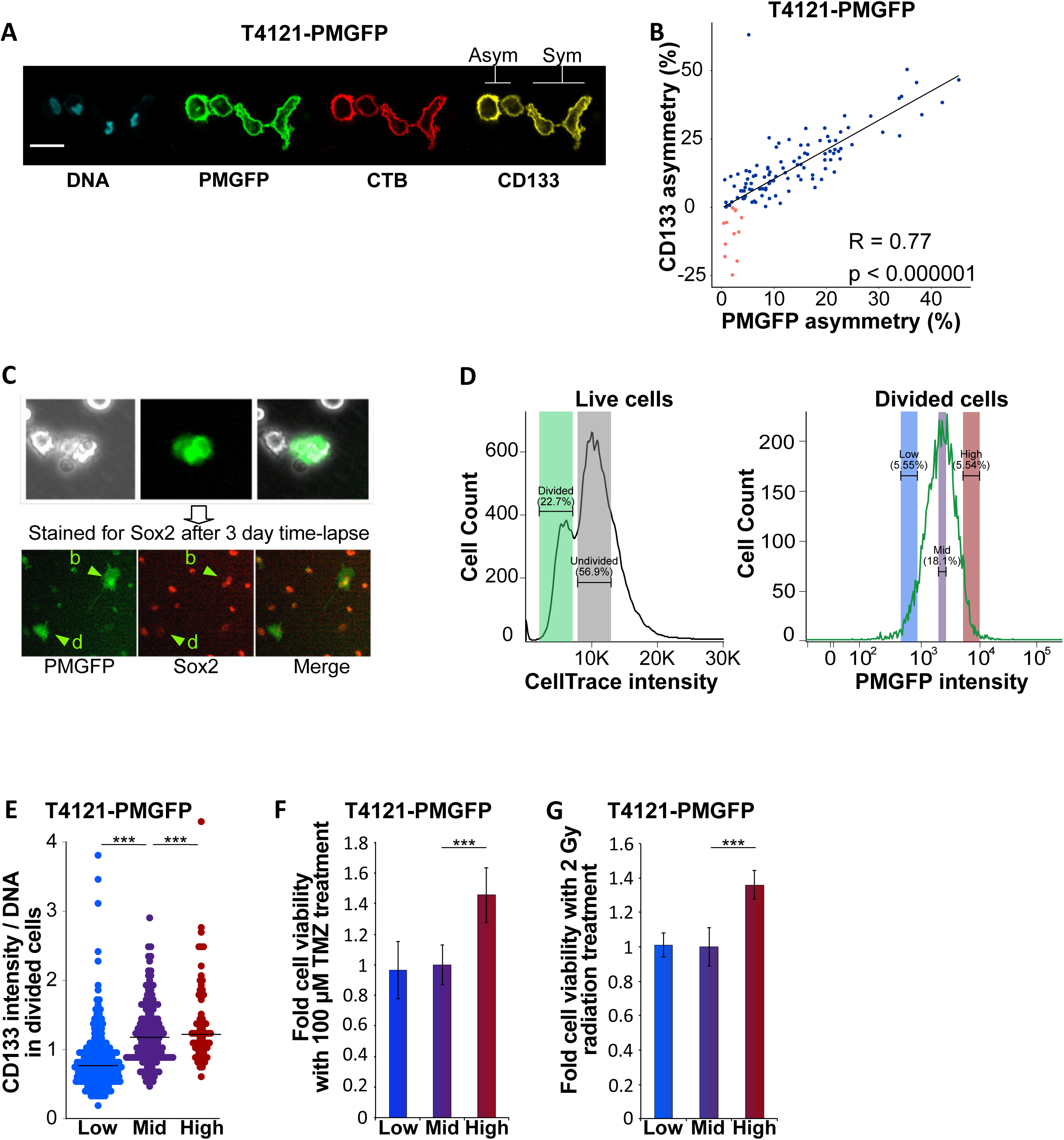
A plasma membrane green fluorescence protein PMGFP reporter system allows the reliable evaluation of cell division mode and reveals functional differences in asymmetrically divided cells. **A** - Confocal microscopy images of two dividing cells in late telophase. Scale bar = 20 µm. DNA staining with Hoechst 333342 (blue). T4121-PMGFP cancer stem cells (CSCs) express the PMGFP construct that localizes in lipid rafts (green). Cholera toxin B (CTB, red) was used as a marker of lipid rafts. CD133 surface staining is shown in yellow. The cell pair on the left exhibits asymmetric (Asym) distribution of PMGFP, CTB and CD133, with one cell receiving more of those markers (left daughter cell). The cell pair on the right exhibits symmetrical (Sym) distribution of the markers. **B** - Quantification of asymmetry percentage during late telophase reveals a correlation between asymmetry of PMGFP and CD133. Each dot represents one cell division. Divisions where PMGFP and CD133 co-segregate on the same daughter cell are marked in blue. Divisions that exhibit segregation of these markers on different daughter cells are marked in red. **C** - Representative images from time-lapse analysis of T4121-PMGFP CSCs. Cells were allowed to divide, fixed 12-72 hr later, and then stained for SOX2. Asymmetry in retrospective PMGFP signal intensity was quantified on both daughter cells: darker cell (d) and brighter cell (b), at the time of mitosis. Divided cells were traced, and the SOX2 levels of the daughter cells were quantified. **D** - FACS analysis of cells after synchronization and mitosis. Once-divided cells exhibited a CellTrace signal intensity that was half (50%) the value of the non-divided cells. Divided cells were gated and then sorted based on their PMGFP signal. Asymmetric divisions constituted 10- 15% of divisions in T4121-PMGFP cells. Hence, the top and bottom 5% of PMGFP cells (PMGFP-high and PMGFP-low) were sorted as asymmetrically divided, and the middle fraction of the PMGFP distribution (PMGFP-mid) was selected as cells that underwent symmetric division. **E** - Immunofluorescence staining quantification of CD133 expression in sorted symmetrically and asymmetrically divided cells. CD133 signal intensity per cell was normalized to DNA intensity per cell. PMGFP-low, PMGFP-mid, and PMGFP-high populations all had a significantly different CD133 expression level, with the highest CD133 level in PMGFP-high and the lowest in PMGFP-low. *** = p<0.000001. **F** – Cell viability of PMGFP-low, PMGFP-mid and PMGFP-high populations after 3-day exposure to 100 µM temozolomide (TMZ). PMGFP-high cells had a significantly higher (*** = p<0.000001) viability. The data represent an average of two biological replicates. **G** - Cell viability of PMGFP-low, PMGFP-mid and PMGFP-high populations 3 days after irradiation with 2 Gy. PMGFP-high cells had a significantly higher (*** = p<0.000001) viability.

### ACD generates progeny with enhanced therapeutic resistance

CSCs are resistant to conventional therapies. To investigate whether the mode of cell division alters therapeutic resistance of the resulting CSC progeny, we isolated dividing daughter cells generated through symmetric or asymmetric cell division using a fluorescence-activated cell sorting (FACS)-based approach (**Supplemental Fig. 1C**). To achieve this, PMGFP CSCs were synchronized in S phase and labeled with CellTrace dye. The cells of uniform PMGFP and CellTrace intensity were enriched by the first round of FACS sorting. The cells were then released from the S phase arrest and 15 hours later gated based on CellTrace intensity to capture the once-divided cells. At the time of release from S phase arrest the cells were subjected to a differentiation-inducing condition that induced ACD in up to 10-15% of the total divisions based on our previous observations (Lathia et al., 2011). Therefore, collecting the top and the bottom 5% of PMGFP intensity levels of the once-divided population likely captured the progeny of ACDs, and the cells with mid PMGFP levels were likely to be progeny of symmetrically divided CSCs (**Fig. 1D**). The fidelity of this approach was confirmed by CD133 staining of sorted populations that revealed the highest levels of CD133 in PMGFP-high cells and lowest CD133 levels in PMGFP-low cells (**Fig. 1E**). This finding is in accordance with co-segregation of CD133 and PMGFP during mitosis (**Fig. 1 A, B**). When challenged with GBM standard-of-care therapies, the progeny with the highest levels of PMGFP inheritance had increased survival after treatment with temozolomide (**Fig. 1F**) and ionizing radiation (**Fig. 1G**). Taken together, these data demonstrate that ACD generates a population of CSC progeny with an enhanced ability to resist conventional therapies.

### ACD co-enriches EGFR and p75NTR, which promotes self-renewal

Lipids rafts concentrate not only CD133, a CSC marker, but also many other signaling molecules, including growth factor receptors that are responsible for therapeutic resistance and tumor growth (Simons and Toomre, 2000). We hypothesized that asymmetric inheritance of lipid rafts results in enrichment of growth factor receptors in the favored daughter cell. We determined the mitotic asymmetry of EGFR and p75NTR and compared it to that of the PMGFP ACD reporter. We chose these receptors based on their important roles in GBM biology. EGFR is a major driver of malignancy (Thorne et al., 2016), while p75NTR facilitates cell infiltration and its ligand is implicated in GBM progression (Johnston et al., 2007; Wang et al., 2018). We found that both EGFR and p75NTR were co-segregated with the PMGFP ACD reporter (**Fig. 2A, B**), confirming previous reports that EGFR can be asymmetrically distributed in GBM CSCs (Cusulin et al., 2015; Sugiarto et al., 2011). Furthermore, co-staining of EGFR and p75NTR demonstrated that these receptors were most often co-enriched in one of the daughter cells during ACD (**Fig. 2A, C**). To verify the co-segregation of our PMGFP reporter and these two growth factor receptors, we utilized our FACS-based approach to isolate progeny generated through symmetric and asymmetric cell divisions (**Fig. 1D, Supplemental Fig. 1C**). We stained populations with low, mid, and high levels of PMGFP for EGFR (**Fig. 2D**) and p75NTR (**Fig. 2E**) and observed that expression levels of these growth factors receptors correlated with PMGFP intensity. To validate these observations in other GBM CSC specimens, we marked lipid rafts with fluorescently labeled CTB and immunostained for EGFR and p75NTR. We detected co-segregation of lipid rafts and these receptors in CSCs from four additional specimens (**Fig. 2F, G**). We were also able to demonstrate the occurrence of asymmetric EGFR segregation in human GBM specimens (**Supplemental Fig. 2A, B**) as well as co-segregation of EGFR with PMGFP in mitotic cells in T4121-PMGFP intracranial xenografts (**Supplemental Fig. 2C**). These data demonstrate co-segregation of our lipid raft reporter with two key growth factors receptors on the same daughter cell during ACD.

**Figure 2.**
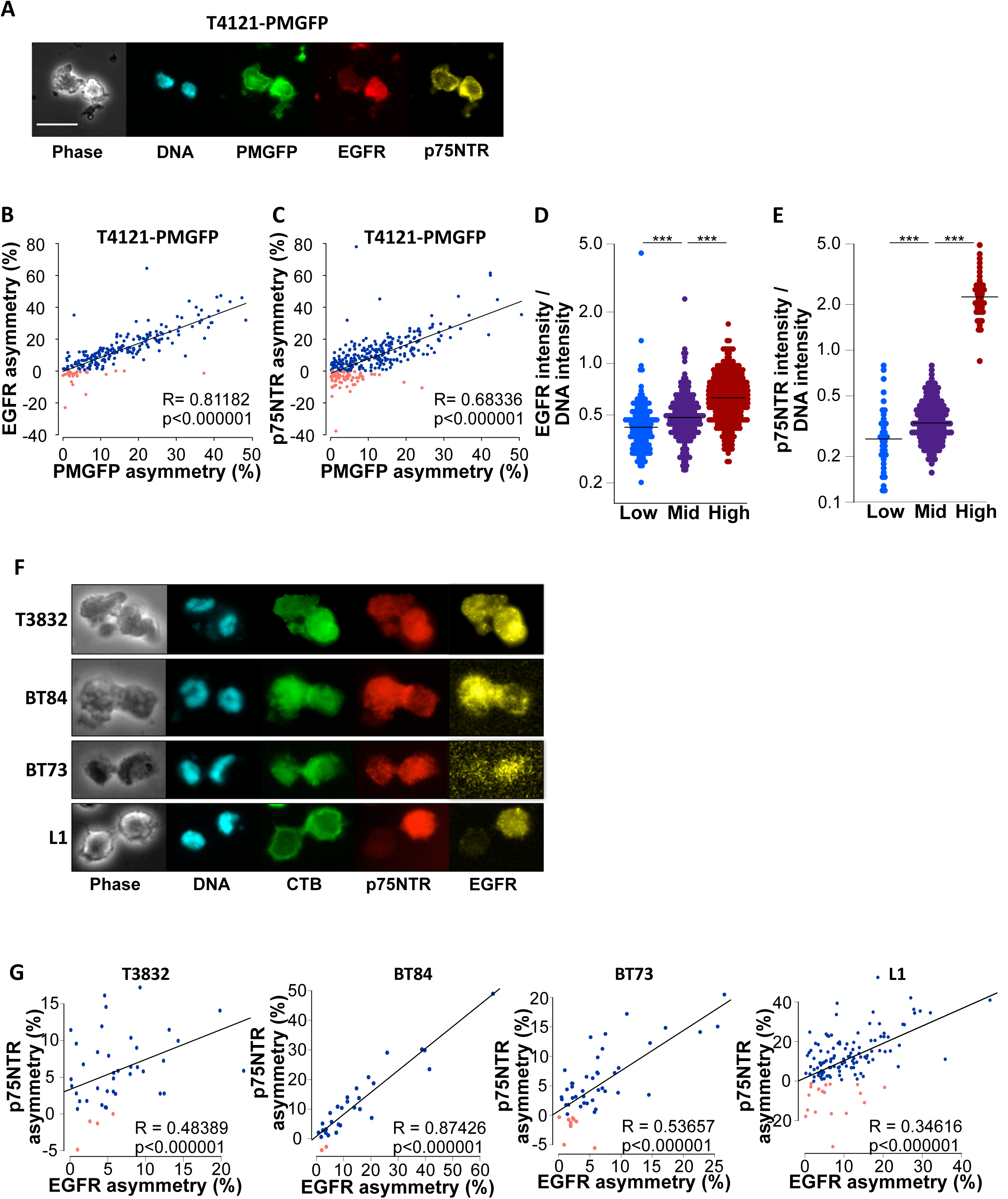
EGFR and p75NTR co-segregate during asymmetric cell division. **A** - Immunofluorescence staining of an asymmetrically divided T4121-PMGFP cell in late telophase. PMGFP (green), EGFR (red), p75NTR (yellow) are shown, and DNA is stained with Hoechst 333342 (blue). **B, C** - Quantification of percentage of asymmetry during late telophase reveals a correlation between the asymmetry of PMGFP and EGFR (**B**) and PMGFP and p75NTR (**C**). Each dot represents one cell division. Divisions where PMGFP and EGFR/p75NTR co-segregated on the same daughter cell are marked in blue. Divisions that exhibited segregation of these markers on different daughter cells are marked in red. **D, E** - Immunofluorescence staining quantification of EGFR (**D**) and p75NTR (**E**) expression in sorted symmetrically and asymmetrically divided cells. EGFR/p75NTR signal intensity per cell was normalized to DNA intensity per cell. PMGFP-low, PMGFP-mid and PMGFP-high populations all had a significantly different EGFR and p75NTR expression level, with the highest expression level in PMGFP-high and the lowest in PMGFP-low. *** = p<0.000001. **F** - Immunofluorescence staining of asymmetrically dividing cells in late telophase from four different glioma stem cell specimens. CTB, used as lipid raft marker (green); p75NTR (red); and EGFR (yellow) are shown; DNA was stained with Hoechst 333342 (blue). **G** - Quantification of asymmetry percentage during late telophase reveals a correlation between asymmetry of EGFR and p75NTR in four different non-PMGFP-expressing glioma stem cell specimens. Divisions where EGFR and p75NTR co-segregated on the same daughter cell are marked in blue. Divisions that exhibited segregation of these markers on different daughter cells are marked in red.

As ACD enriches EGFR and p75NTR on the same daughter cell, we next assessed the biological importance and interaction of the signaling activities of the two receptors in CSC maintenance. We subjected CSCs to a serum-based differentiation paradigm (Patel et al., 2014), which reduced the self-renewal capacity of the cells (**Fig. 3A, B**). Stimulation of both receptors under this differentiation condition restored self-renewal capacity (**Fig. 3B**), but activation of each receptor alone was not sufficient to override the differentiation-inducing effects of serum (**Fig. 3C)**. This observation suggests that the activity of these two receptors co-inherited during ACD cooperates to enhance self-renewal capacity of one of the daughter cells. As expected, knockdown of p75NTR attenuated the ability of EGF and p75NTR ligands to override the effect of serum to suppress self-renewal (**Fig. 3D, E**), validating that the two signaling pathways synergize to maintain the CSC phenotype.

**Figure 3.**
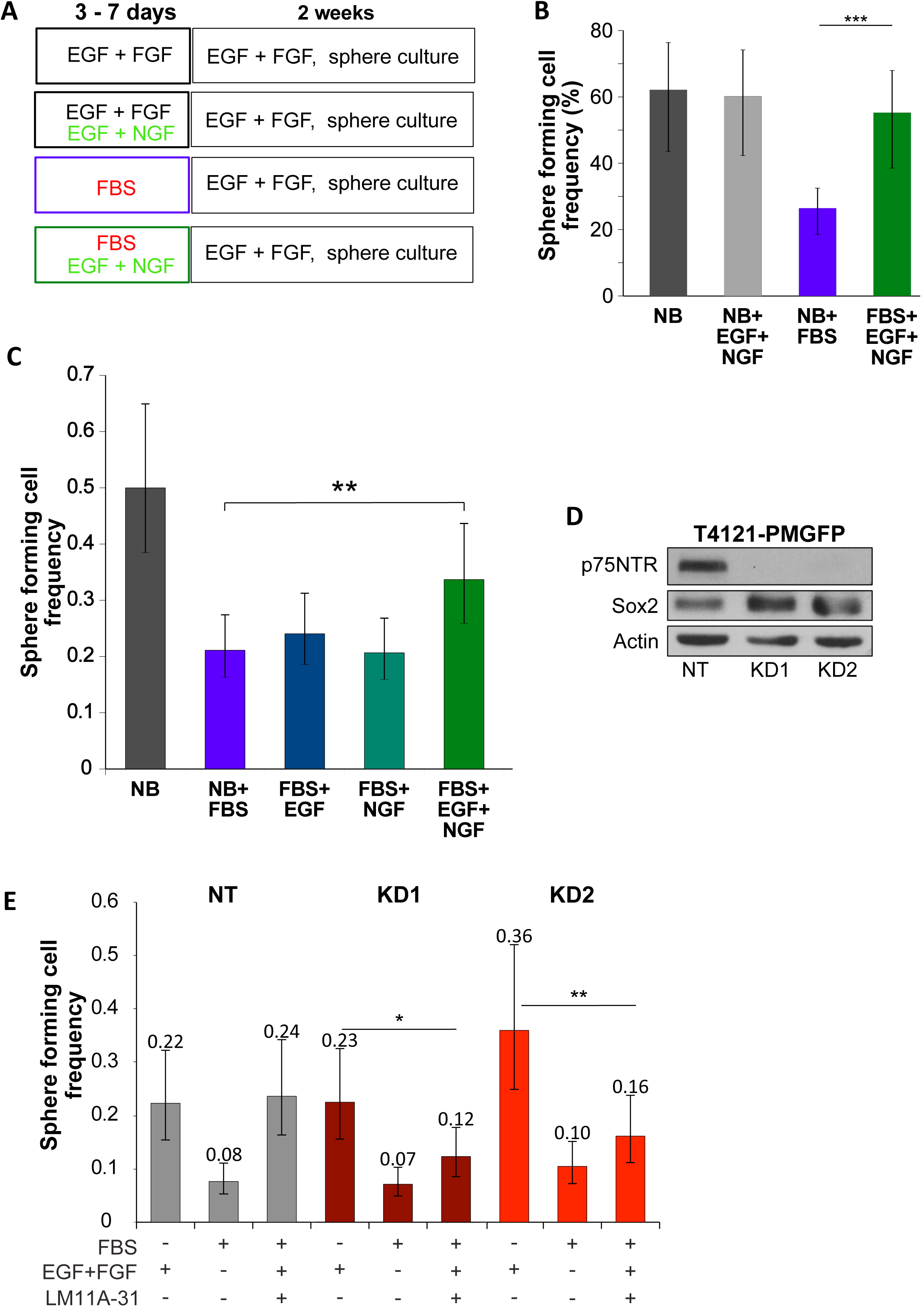
Alteration of the p75NTR axis modifies CSC phenotypes after differentiation. **A** - Experimental design for the differentiation experiment. T4121-PMGFP CSCs were subjected to 3 days of pretreatment with either CSC medium (containing epidermal growth factor (EGF) and fibroblast growth factor 2 (FGF2)), CSC medium with the addition of nerve growth factor (NGF), CSC medium with the addition of 10% FBS without EGF and FGF, or CSC medium with the addition of 10% FBS and EGF and NGF. The cells were then plated for limiting-dilution assay in CSC medium and assessed for self-renewal 2 weeks later. **B** - Self-renewal capacity of T4121-PMGFP cells after differentiation with or without NGF stimulation. *** p<0.000001. **C** - Self-renewal capacity of T4121-PMGFP cells after differentiation using 10% FBS with and without growth factor stimulation (EGF, NGF, or combination). ** p=0.0144. **D** - Immunoblotting showing expression of p75NTR and SOX2 in T4121-PMGFP cells expressing p75NTR knockdown shRNAs (KD1 and KD2) compared to non-targeting (NT) shRNA. **E** - Self-renewal capacity of T4121-PMGFP cells in stem cell medium (NB) alone or in the presence of the p75NTR ligand LM11A-31 (NB+LM, 100 nM), 10% FBS, or FBS with LM11A-31 together (FBS+LM). Non-targeting shRNA-containing cells (NT) were compared to knockdown shRNA (KD1 and KD2)-containing cells. *** p<0.000001.

### p75NTR ligands rescue EGFR inhibition

EGFR signaling is activated in the majority of GBM cases making this receptor a candidate for therapeutic target (Halatsch et al., 2006; Voelzke et al., 2008). EGFR targeting through its kinase inhibition, however, failed to show therapeutic benefit in clinical trials (Thorne et al., 2016). Based on our current observation of EGFR and p75NTR co-segregation and their reportedly overlapping downstream signaling pathways (Longo and Massa, 2013), we hypothesized that signaling activity from p75NTR compensates for EGFR inhibition. To test this hypothesis, we stimulated cells with p75NTR ligands in the presence of erlotinib, which inhibits EGFR kinase activity. Erlotinib suppressed autophosphorylation of full-length and truncated mutant EGFR as well as SOX2 expression (**Fig. 4A**). The reduction in both autophosphorylation of EGFR Y1086 and SOX2 expression was rescued when cells were stimulated with natural ligands of p75NTR expressed in the brain, nerve growth factor (NGF) and brain-derived neurotrophic factor (BDNF; **Fig. 4A**). We assessed known downstream signaling nodes of these receptors and found that the erlotinib-mediated reduction in STAT3 and AKT phosphorylation was overridden by p75NTR natural and synthetic ligands. This effect was not as pronounced for another known downstream signaling mediator, ERK (**Fig. 4B**). Importantly, STAT3 is a well-established CSC maintenance signaling node (Ganguly et al., 2018) that is activated through EGFR signaling (Sartor et al., 1997). Our results indicate that EGFR inhibition is not sufficient to suppress STAT3 activity when converging p75NTR signaling is activated.

**Figure 4.**
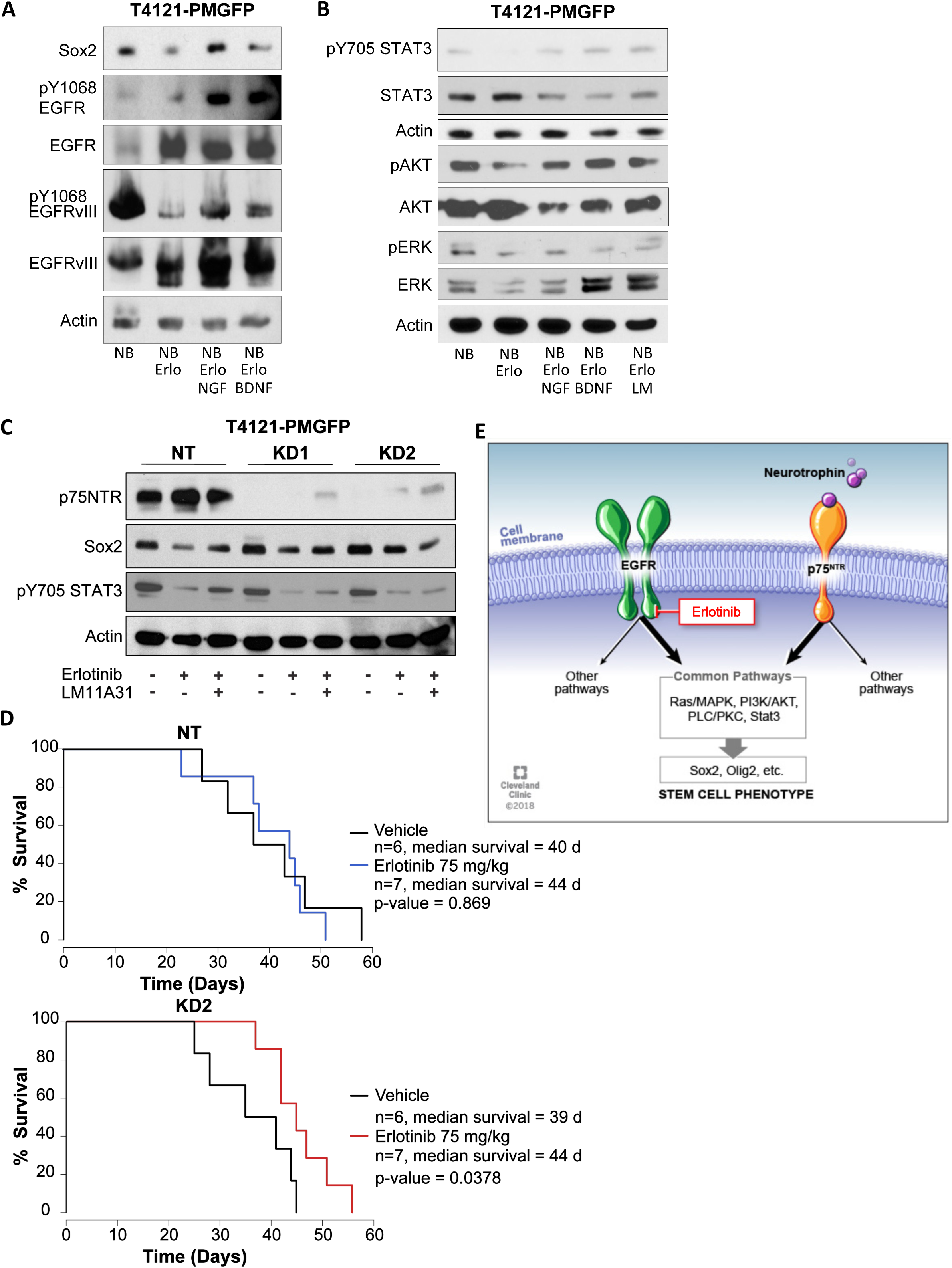
Alteration of the p75NTR axis modifies response to EGFR inhibition. **A** - Immunoblotting for SOX2 and EGFR receptor activation in T4121-PMGFP cells after a 3-day treatment with stem cell medium (NB), the EGFR inhibitor erlotinib (Erlo; 3 µM) or a combination of erlotinib and NGF (Erlo+NGF; 100 nM) or BDNF (Erlo+BDNF; 100 nM) to stimulate the p75NTR receptor axis. **B** - Immunoblotting for STAT3, AKT and ERK activation in T4121-PMGFP cells after a 3-day treatment with stem cell medium (NB), the EGFR inhibitor erlotinib (NB+Erlo; 3 µM) or a combination of erlotinib and NGF (NB+Erlo+NGF; 100 nM), BDNF (NB+Erlo+BDNF; 100 nM) or the p75NTR ligand LM11A-31 (NB+Erlo+LM; 100 nM) to stimulate the p75NTR receptor axis. **C** - Immunoblotting for EGFR receptor activation and SOX2 expression in non-targeting (NT) and knockdown (KD1 and KD2) T4121-PMGFP glioma stem cells. Cells were treated for 3 days with CSC medium (NB) alone, CSC medium with the addition of the EGFR inhibitor erlotinib (Erlo; 3 µM) or erlotinib with the p75NTR ligand LM11A-31 (Erlo+LM; 100 nM). **D** - Survival of animals intracranially implanted with T4121-PMGFP cells containing non-targeting (NT) shRNA or p75NTR knockdown (KD1) shRNA. Animals in the vehicle group were treated with 0.05% methylcellulose solution. The erlotinib group received 75 mg/kg per day. n = number of animals in each group. Median survival and p-value as determined by log-rank test comparing the vehicle and erlotinib groups are shown. **E** - Schematic depicting the proposed model of signaling that occurs in GBM CSCs upon EGFR and p75NTR co-segregation during ACD. The two receptors signal through similar signaling pathways that promote the stem cell phenotype. Upon inhibition of EGFR using erlotinib, p75NTR takes over the downstream stimulation to maintain the stem cell program.

### p75NTR attenuates the therapeutic efficacy of EGFR inhibition

To further understand the role of the p75NTR receptor in the context of EGFR targeted therapy, we next treated p75NTR knockdown cells with erlotinib to suppress EGFR kinase activity. While the reduction in p75NTR did not dramatically suppress SOX2 expression (**Fig. 4C, D**) or self-renewal capability of CSCs (**Fig. 3E)**, knockdown of this receptor attenuated the ability of p75NTR ligand to rescue SOX2 expression suppressed by erlotinib (**Fig. 4C, D**). To determine whether p75NTR knockdown increased the sensitivity of CSCs to erlotinib, we assessed IC_50_ values and found that the IC_50_ was lower in cells where p75NTR was knocked down using the KD2 shRNA construct (**Supplemental Fig. 3A**). Using this p75NTR knockdown CSC population, we tested the importance of p75NTR in resistance to erlotinib using an orthotopic mouse xenograft model. We first assessed the consequence of p75NTR knockdown on tumor growth and found no significant difference in survival between the mice injected with CSCs expressing a non-targeting shRNA control and those expressing p75NTR shRNA (**Supplemental Fig. 3B**). We next determined the minimal effective dose of erlotinib in vivo by evaluating a dose rage from 5 to 100 mg/kg, similar to a previously reported range (Sarkaria et al., 2006). We found that 100 mg/kg significantly increased the survival of tumor-bearing mice, while lower doses did not increase survival (**Supplemental Fig. 3C**). We reasoned that if p75NTR compensates for EGFR function that is suppressed by erlotinib, xenograft tumors originated from p75NTR knockdown CSCs would become susceptible to erlotinib at a dosage that did not affect the tumorigenicity of control CSCs. Indeed, we found that 75 mg/kg erlotinib increased the survival of mice bearing p75NTR knockdown tumors but not the survival of mice with control tumors (**Fig. 4E**). These results demonstrate that the p75NTR signaling axis compensates for EGFR signaling to override the therapeutic efficacy of EGFR inhibition.

### Discussion of key findings

ACD is an essential cell division mode that enables simultaneous maintenance of a stem cell population and the generation of differentiated progeny during embryogenesis, organogenesis, tissue homeostasis, and tissue regeneration (Cicalese et al., 2009; Knoblich, 2008; Venkei and Yamashita, 2018). ACD has been reported in many advanced cancers, but the importance of ACD during tumorigenesis has yet to be fully elucidated. Technical challenges such as difficulty tracking the fate of daughter cells after different modes of cell division and/or the lack of lineage-tracing systems have prevented studies from elucidating the contribution of ACD to tumorigenic processes. While cell fate tracking has demonstrated dynamic evolution in cancer and therapeutic response, this has not been fully linked to cell fate choice. Moreover, in many studies, ACD is often defined retrospectively based on two progeny with different phenotypes, and this type of retrospective view hampers prospective mechanistic analysis (Chen et al., 2014; Lathia et al., 2011). The current work represents a new opportunity to investigate ACD in a prospective manner by virtue of a fluorescent protein-based reporter of asymmetric mitotic inheritance of key signaling molecules. This reporter system enables the previously unfeasible investigation of the impact of ACD on cell fate decisions as well as the FACS machine-based collection of the large cell numbers required for molecular and phenotypic characterization of ACD progeny.

Previous work from our group and others has shown that ACD is not the dominant mode of cell division used by CSCs during tumor growth (Lathia et al., 2011; Xin et al., 2012). These findings have also been confirmed using mathematical modelling to suggest an evolutionary disadvantage for ACD (Daynac et al., 2018; Guichet et al., 2016; Mukherjee et al., 2016; Obernier et al., 2018; Sugiarto et al., 2011; Tomasetti and Levy, 2010; Tominaga et al., 2019). However, these assessments have not been done in the context of the selective pressures induced by therapies. Our current findings demonstrate that ACD becomes advantageous during therapeutic stress by generating a daughter cell with enhanced capacity to withstand therapies. ACD would likely result in an overall decrease in growth but helps to preserve a population of cells with an evolutionary advantage that subsequently drives tumor recurrence. This paradigm may be useful when designing and assessing the efficacy of pathway-specific inhibitors, such as those targeting EGFR activity, which may require concomitant neutralization of other signaling pathways to achieve a durable therapeutic response.

Tumors consist of heterogeneous populations of tumor cells, and this cellular heterogeneity has been implicated in tumor therapeutic resistance (Osuka and Van Meir, 2017; Qazi et al., 2017; Richardson and Siemann, 1997). The net fitness of the tumor cannot be determined solely by that of tumorigenic cells such as CSCs or the most proliferative cells in the tumor; a dynamic interaction among the different types of tumor cells and their interaction with stromal cells are also critical factors. Furthermore, reciprocal crosstalk between CSCs and more differentiated tumor cells may contribute to tumor growth (Silver and Lathia, 2018; Wang et al., 2018). In this context, ACD may contribute to overall tumor growth by generating heterogeneous populations of cells that form a mutually beneficial paracrine network involving BDNF, a p75NTR ligand. Analysis of cell division mode should also be expanded to non-stem cancer cells that can revert to a CSC phenotype as a result of chemotherapy or microenvironment-induced stresses.

While these studies provide insight into the role of ACD in therapeutic resistance and CSC fate choice, the underlying fundamental molecular mechanisms have yet to be determined. Long non-coding RNAs as well as transcription factors (Liu et al., 2018; Wang et al., 2016) have been demonstrated to be asymmetrically distributed in CSCs, which may be a potential upstream mechanism of ACD. Uncovering the molecular mechanism driving ACD represents a priority for future studies (Zimdahl et al., 2014). Asymmetrically generated progeny showed differential sensitivity to TMZ and radiation, standard-of-care therapeutics for GBM. Further studies are required to establish the molecular mechanism downstream of ACD that provides therapeutic resistance. Such studies will reveal exploitable targets to enhance the efficacy of conventional treatments. The cell division reporter and sorting strategies we describe here provide a critical platform to perform such studies.

## Materials and Methods

### Xenograft maintenance

Established GBM xenografts (T4121, T3832, BT84, BT73 and L1) were previously reported (Bao et al., 2006; Cusulin et al., 2015; Schonberg et al., 2015; Srinivasan et al., 2016) and were obtained via a material transfer agreement from Duke University, University of Florida, and the University of Calgary, where they were originally established under IRB-approved protocols that facilitated the generation of xenografts in a de-identified manner from excess tissue taken from consented patients. For experimental studies, GBM cells were dissociated from established xenografts under Cleveland Clinic-approved Institutional Animal Care and Use Committee protocols. Xenografts were passaged in immune-deficient NOD.*Cg-Prkdc*^*scidIl2rgtm1Wjl*^/SzJ (NSG) mice (obtained from The Jackson Laboratory, Bar Harbor, ME, USA) to maintain tumor heterogeneity. Six-week-old female mice were unilaterally injected subcutaneously in the flank with freshly dissociated human GBM cells, and animals were sacrificed by CO_2_ asphyxiation and secondary cervical dislocation when tumor volume exceeded 5% of the animal’s body weight.

### CSC isolation

Xenografted tumors were dissected and mechanically dissociated using papain dissociation kits (Worthington Biochemical Corporation, Lakewood, NJ, USA), and cells were cultured overnight in neurobasal medium (Life Technologies, Carlsbad, CA, USA) supplemented with B27 (Life Technologies), 1% penicillin/streptomycin (Life Technologies), 1 mM sodium pyruvate, 2 mM L-glutamine, 20 ng/mL EGF (R&D Systems, Minneapolis, MN, USA), and 20 ng/mL FGF-2 (R&D Systems) in a humidified incubator with 5% CO_2_. CSCs were enriched using the CD133 Magnetic Bead Kit for Hematopoietic Cells (CD133/2; Miltenyi Biotech, San Diego, CA, USA) and cultured in supplemented neurobasal medium. This enrichment method reliably enriches CSCs that have increased self-renewal compared with their non-CSC counterparts (Bao et al., 2006; Schonberg et al., 2015). Cells were cultured in supplemented neurobasal medium until the day they were used.

### Intracranial cell injection and erlotinib treatment

Five-to-eight-week old NSG mice were anesthetized using isoflurane and positioned for intracranial injection using a stereotaxic frame (Kopf Instruments, Tujunga, CA, USA). Five microliters of a single cell suspension of GBM CSCs was injected into the left striatum at a concentration of 10,000 cells/animal. Two weeks after injection, animals were randomized into treatment and control groups. Daily gavage with 100 µL of either 0.5% methylcellulose (vehicle group) or a suspension of erlotinib in 0.5% methylcellulose (erlotinib group) was performed for 4 weeks. Animals were monitored and euthanized when neurological symptoms developed. For the experiments in Supplemental Figure 3B and C, female mice were used. For the experiments in Figure 4E, male mice were used.

### PMGFP expression in CSCs

A PMGFP plasmid from Addgene (plasmid #21213) was linearized by digestion with the restriction enzyme NruI and transfected into T4121 CSCs using lipofectamine. A stable resistant population was selected with G418 (1 mg/ml) and sorted using FACS to enrich for GFP-positive cells. A PMGFP-expressing stable population was maintained in medium containing a reduced amount of G418 (0.3 mg/ml).

### Mitotic shake-off

To analyze protein expression on daughter cells at the time of mitosis, we enriched for mitotic cells using mitotic shake-off. GBM CSCs were cultured adherently as a monolayer on Geltrex-coated plates. The cells were synchronized in S-phase using the addition of 2 mM thymidine to the medium for 12 hours. After synchronization, the cells were released into thymidine-free medium for 12-15 hours to allow the cells to progress through the cell cycle and reach mitosis. At this point, the plates were subjected to gentle vortexing to allow the rounded-up mitotic cells to detach from the plates. The detached cells were washed from the plate and centrifuged onto poly-lysine-coated coverslips. The cells were then promptly fixed with 4% paraformaldehyde and stained.

### Time-lapse fate-decision tracing

GBM CSCs (T4121) and CSCs expressing PMGFP (T4121-PMGFP) were plated adherently onto Geltrex-coated 6-well plates as a mixed culture at a ratio 1:5, with a total of 200,000 cells/well. The cells were synchronized in S-phase using the addition of 2 mM thymidine to the media for 12 hours. The cells were then released into CSC medium with 10% fetal bovine serum (FBS) but without EGF/FGF to increase the rate of ACD (based on previously published data). Using a sterile needle, a straight 0.5 cm scratch was made at the bottom center of each well. Time-lapse microscopy was then initiated using a Leica CTR6500 time-lapse microscope with Tempcontrol Digital set-up at 37°C in humidified air with 5% CO2, with phase images taken every 5 min and green fluorescence captured every 30 min to avoid phototoxicity and bleaching. Acquisition of images was performed using Leica LAS X Life Science software. For each well, 6 fields of view adjacent to the scratch were captured. After 72 hours, the time-lapse was stopped, and the cells were promptly fixed using 4% paraformaldehyde in PBS and subjected to immunofluorescence staining for SOX2. The staining was then reviewed using a Leica DM5000B microscope equipped with a Leica DFC310 FX Digital Color Camera. Images from time-lapse microscopy were exported as .tiff format and analyzed using NIH ImageJ. The exact locations and the individual cells captured by time-lapse imaging were identified using the scratches on the bottom of the wells as reference landmarks. Digital images of the staining were quantified to determine the levels of SOX2 expression in each progeny and correlated with the brightness of PMGFP that each daughter cell received during mitosis.

### FACS-based isolation of divided cells

To enrich for mitotic cells, we synchronized GBM CSCs (approximately 100 million cells) cultured as spheres at G1/S-phase border by adding 2 mM thymidine-containing CSC medium for 12 hours. The spheres were then dissociated into single cells using Accutase (Life Technologies, Carlsbad, CA), counted, and labeled with CellTrace Far Red dye (Life Technologies, Carlsbad, CA) by suspending the cells in 40 mL of media with 25 µL of CellTrace dye per 100 million cells for 20 min at 37°C. The cells were then pelleted and released into CSC medium without thymidine for 6 hours to let them progress to mid-S phase of the cell cycle. The cells were then suspended in FACS buffer (CO_2_-independent medium (Life Technologies) + B27) containing 2 mM thymidine to prevent further progress through the S phase and subjected to FACS at 4°C over the next 6 hours to isolate a homogeneous population in terms of green and far red intensity. Subsequently, the sorted cells were simultaneously released into Neurobasal medium containing 10% FBS and B27 without thymidine to increase the rate of ACD (based on previously published data). After 15 hours, a significant percentage of cells had undergone one mitosis, at which point the cells were subjected to a second FACS: live cells were gated based on CellTrace Far Red intensity to collect only the cells that had divided once (Fig. 1D) as well as gated for the bottom and top 5% to collect asymmetrically divided cells and for a narrow range in average PMGFP intensity to select for symmetrically divided daughter cells. Isolated cells populations were further subjected to immunofluorescence staining and functional proliferation assays. For immunofluorescence staining, the cells were centrifuged onto Geltrex-coated cover slips and incubated for 1-2 hours to allow the cells to attach prior to fixing.

### Immunofluorescence staining

Cells for immunofluorescence staining were fixed on cover slips in 6-well plates using 4% paraformaldehyde in PBS for 15 min, followed by rinsing with PBS. Fixed cells were blocked with 2% donkey serum (EMD Millipore, Carlsbad, CA) for 1 hour. For EGFR and p75NTR staining, we permeabilized cells with 0.01% Triton added to the blocking solution. The samples were then incubated overnight at 4°C with primary antibodies against CD133, EGFR and/or p75NTR; washed three times with PBS; and incubated for 1 hour at room temperature with secondary antibodies: DyLight-649-conjugated donkey anti-rabbit IgG, Cy3-conjugated donkey anti-mouse IgG and DyLight-649-conjugated donkey anti-mouse IgG (Jackson Immunoresearch). The samples were washed three times with PBS and incubated for 15 min in PBS containing Hoechst 33342 (100 ng/mL) and/or Alexa-488-or Alexa-594-conjugated CTB. The cover slips were then mounted on slides in gelvatol mounting medium (PVA (Sigma-Aldrich), glycerol (Sigma-Aldrich), sodium azide (Fisher), Tris-Cl pH 8.5) and subjected to fluorescence microscopy using a Leica DM5000B microscope equipped with a Leica DFC310 FX Digital Color Camera. Images were captured at 40x magnification using a dry objective.

### Cell proliferation analysis to determine the effect of therapeutics

To assess the effect of TMZ on proliferation, symmetrically and asymmetrically divided cells derived from GBM CSCs were isolated using the FACS-based approach and plated into 96-well plates at a concentration of 2,000 cells/well in 100 µL of either CSC medium with 100 µM temozolomide or DMSO (1:1000) in CSC medium as a control, with 8 wells per condition. After 3 days of incubation at 37°C with 5% CO_2_, proliferation was assessed using Cell Titer Glo: 100 µL of reagent was added per well and incubated in the dark for 15 min. Luminescence was registered using a Victor 3 multi-well plate reader (PerkinElmer). The experiment was repeated twice.

To analyze the effect of radiation, CSC progeny generated via different modes of cell division were isolated using the FACS-based approach and plated into 96-well plates at a concentration of 2,000 cells/well in 100 µL of CSC medium. The cells were then irradiated with a total dose of 0 or 2 Gy using a Shepherd Cs137 irradiator, with 6 wells per condition. After 3 days of incubation at 37°C with 5% CO_2_, proliferation was assessed using Cell Titer Glo: 100 µL of reagent was added per well and incubated in the dark for 15 min. Luminescence was registered using a Victor 3 multi-well plate reader.

The effect of erlotinib was determined as follows: GBM CSCs were plated in 96-well plates at a concentration of 2,000 cells/well in 100 µL of either CSC medium with 0.3 - 80 µM erlotinib or DMSO (1:1000) in CSC medium as a control, with 3 wells per condition. After 3 days of incubation at 37°C with 5% CO_2_, proliferation was assessed using Cell Titer Glo: 100 µL of reagent was added per well and incubated in the dark for 15 min. Luminescence was registered using a Victor 3 multi-well plate reader.

### Self-renewal assay

To determine the self-renewal capacity of GBM CSCs, 500,000 cells were plated adherently as a monolayer using Geltrex-coated 6-cm tissue culture plates. Cells were cultured under the following conditions for 3 days: NB - CSC medium; NB+EGF+NGF - CSC medium with the addition of extra 20 ng/mL EGF and 100 ng/mL NGF; FBS - neurobasal medium with 10% FBS, L-glutamine, B27, penicillin/streptomycin, and sodium pyruvate; FBS+NGF - FBS medium with 100 nM NGF; LM - CSC medium with 100 nM LM11A-31 (a synthetic p75NTR ligand); FBS+LM - FBS medium with 100 nM LM11A-31; Erlo - CSC medium with the addition of 3 µM erlotinib; Erlo+NGF/BDNF/LM - CSC medium + 3 µM erlotinib + 100 nM NGF/BDNF/LM. After 3 days, cells were dissociated and plated in 96-well suspension plates at 100, 50, 25, 12, 6 and 3 cells per well for limited-dilution analysis in 100 µL of CSC medium. Two plates per condition were used. After 14 days, wells containing spheres were counted, and self-renewal was analyzed using an online tool (http://bioinf.wehi.edu.au/software/elda/)(Hu and Smyth, 2009).

### Immunoblotting

GBM CSCs were collected from adherent monolayer cultures, and whole cell lysates were made in a lysis buffer containing 10% NP-40 (Sigma-Aldrich), 1 mM EDTA, 150 mM NaCl, 10 mM Tris-Cl pH 7.5 supplemented with protease and phosphatase inhibitor cocktails (Sigma-Aldrich). Protein expression was analyzed by immunoblotting for expression of phospho-EGFR (Y1068, 1:1000, Cell Signaling, Danvers, MA, USA), total EGFR (1:1000, E235, Abcam, Cambridge, UK), SOX2 (1:500, R&D Systems), p75NTR (1:1000, Cell Signaling), phospho-STAT3 (1:1000, Y705, Cell Signaling), STAT3 (1:1000, Cell Signaling), pAKT (1:1000, Cell Signaling), AKT (1:1000, Cell Signaling), pMAPK (1:500, Cell Signaling), MAPK (1:1000, Cell Signaling). Anti-beta-actin (1:5000, Santa Cruz Biotechnology, Dallas, TX, USA) was used as a loading control.

### Lentivirus preparation and p75NTR knockdown

Using Biotool DNA transfection reagent (bimake.com, Houston, TX), 293T cells were co-transfected with ps.PAX2, p.MD2.G (Addgene, Cambridge, MA) and lentiviral vectors: non-targeting control (PLK0.1) or p75NTR-targeting MISSION shRNA constructs (Sigma-Aldrich, TRCN0000058153 and TRCN0000058153). The medium was changed 8 hours post-transfection, and viral supernatants were collected 12, 24 and 36 hr later. Viral particles were concentrated using polyethylene glycol precipitation and stored at −80 C. T4121-PMGFP CSCs were infected with the concentrated viral supernatants, selected with 2 mM puromycin for 48 hr. After knockdown was confirmed by immunoblotting, cells were used for further experiments.

### Quantification and statistical analysis

#### Asymmetry quantification

To assess asymmetry in the expression of markers between daughter cells, we developed an ImageJ macro for batch digital image processing of CSCs in late telophase, as described previously (Lathia et al., 2011). The macro automatically measures the fluorescence intensity of manually outlined daughter cells and background fluorescence. To quantify the percent asymmetry between daughter cells, a web-app called Asymmetry was built using R language and the packages shiny, ggplot2, cowplot, dplyr and ggExtra. The app quantified the percent asymmetry of each marker using the following formula:

[(Cell1Intensity-Background1)-(Cell2Intensity-Background2)]X100 / [(Cell1Intensity-Background1)+(Cell2Intensity-Background2)] = %Asymmetry

Pearson’s correlation was calculated to determine the significance of cosegregation between molecules.

#### Immunofluorescence intensity quantification

Using high-throughput automated single-cell imaging analysis (HASCIA), which was described previously (Chumakova et al., 2019), the expression of growth factor receptors on FACS-sorted daughter cells was analyzed. First, the HASCIA image processing script and ImageJ v1.52k were used to obtain single-cell measurements of marker intensity. Then, using the HASCIA web-app, the expression was normalized to DNA intensity, and relative expression difference between groups was assessed using t-test.

#### Statistical analysis

As specified in the text, t-test, Pearson’s correlation or log-rank test were performed to calculate statistical significance, with p values detailed in the text and figure legends. P values < 0.05 were considered significant. Data analysis was done Office Excel 2013 (Microsoft, Redmond, WA), custom R scripts in RStudio using packages survival and limdil, as well as HASCIA.

## RESOURCES

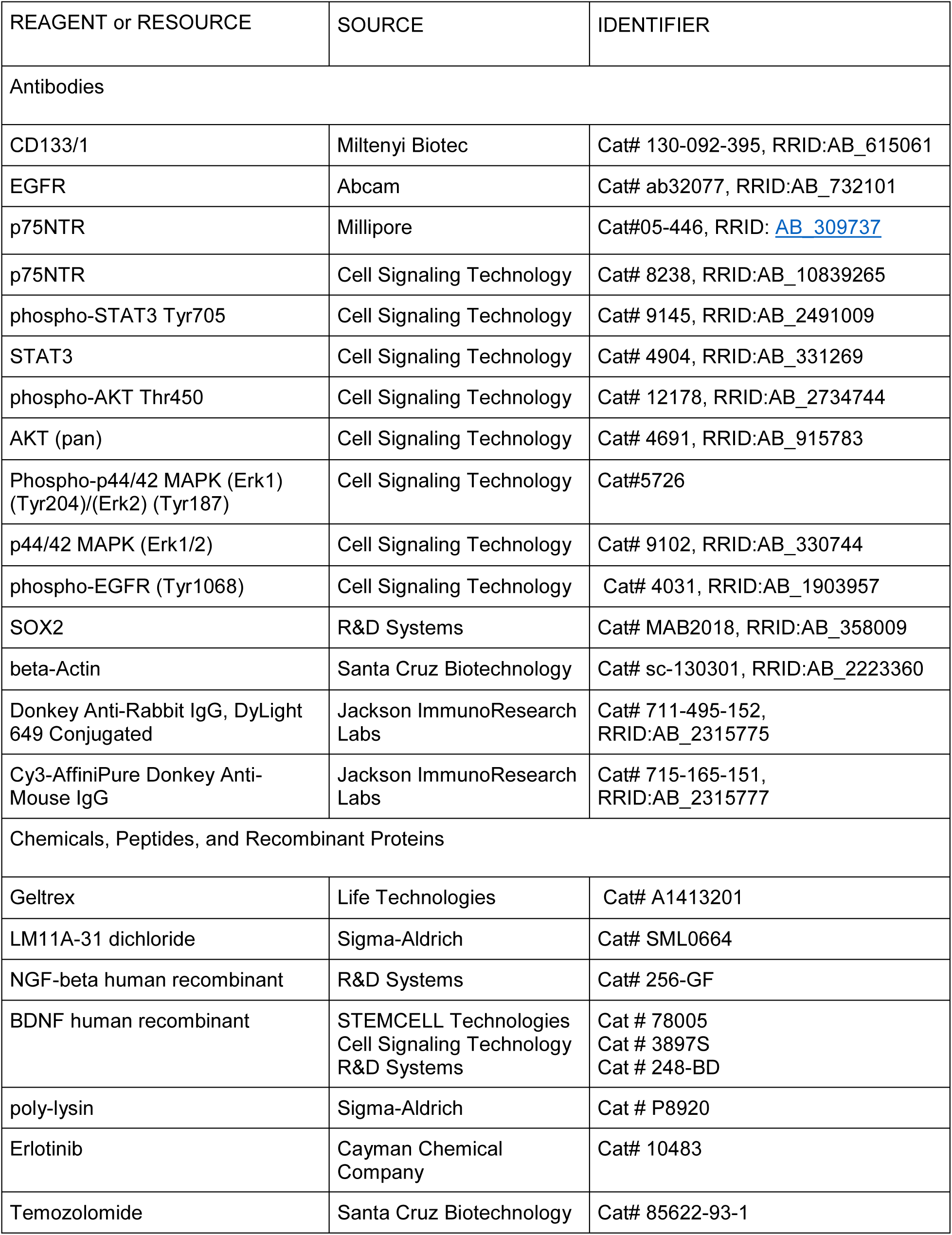
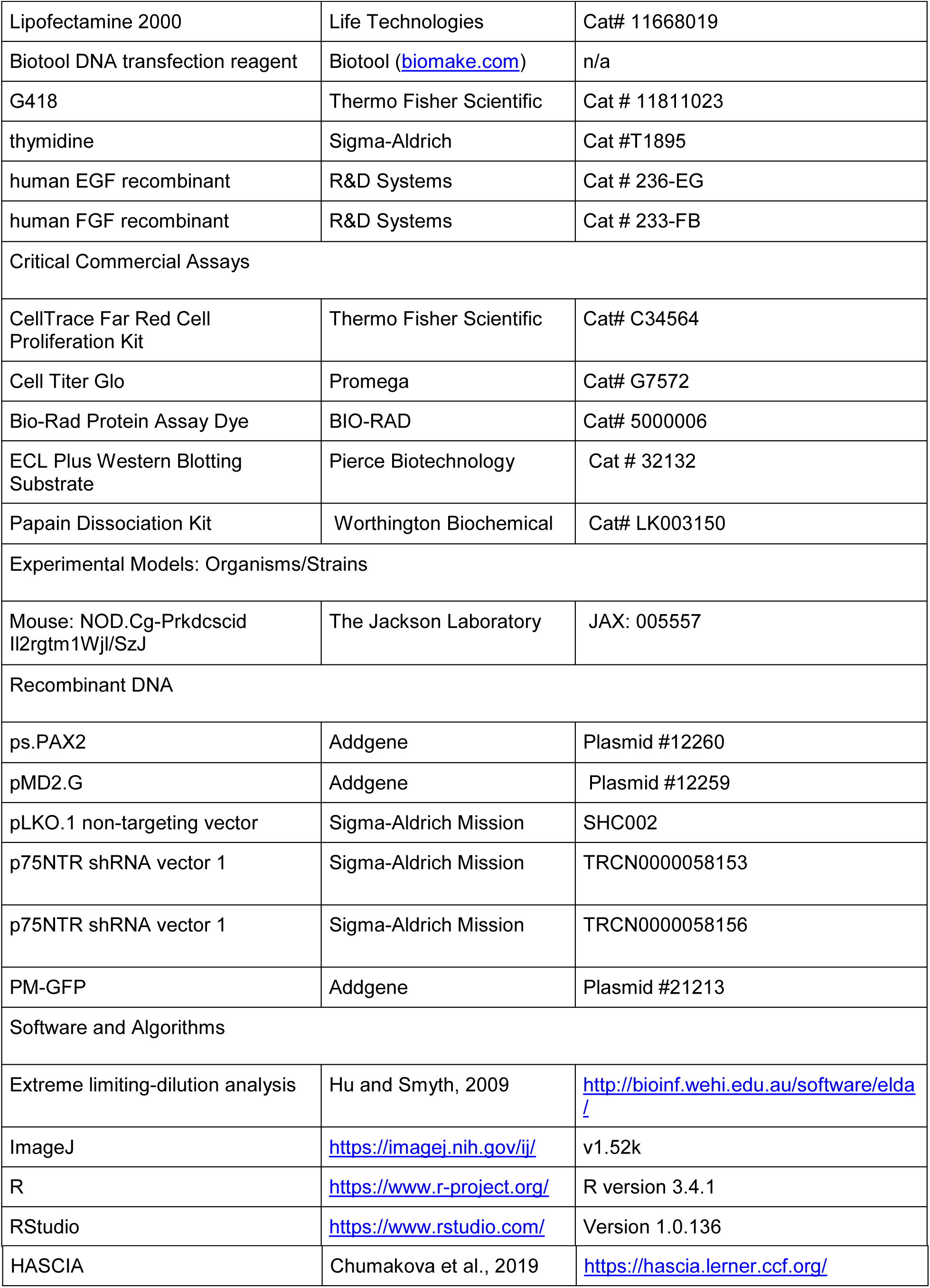

## Acknowledgements

The authors would like to thank the members of the Lathia laboratory and Dr. Rajappa Kenchappa (Mayo Clinic, Jacksonville) for critical feedback and discussions and Amanda Mendelsohn from the Center for Medical Art and Photography at the Cleveland Clinic for her assistance. We thank Drs. Artee Luchman and Sam Weiss at the University of Calgary for providing experimental models. We also would like to thank Upashruti Agrawal, Kayla Pfaff, Peter Jeong, Bridget Corrigan, Brennen Keuchel, Malini Kamini, Vid Yogeswaran, and Helle Wohlleben for their technical assistance. We thank Masahiko Sato and Sachiko Sato from Laboratory of DNA Damage Responses and Bioimaging, CHU de Québec, Canada for their contribution to the concept of time-lapse microscopy analysis. We thank Dr. Erin Mulkearns-Hubert for editorial assistance. These studies were supported by the NIH (R03 CA215939 to M.H.), Cleveland Clinic (J.D.L.), Case Comprehensive Cancer Center (J.D.L.), and the Sontag Foundation (J.D.L). The authors declare no competing financial interests.

## Abbreviations

ACD: asymmetric cell division
BDNF: brain-derived neurotrophic factor
CSC: cancer stem cell
CTB: cholera toxin B
EGFR: epidermal growth factor receptor
GBM: glioblastoma
NGF: nerve growth factor
p75NTR: p75 neurotrophin receptor
PMGFP: plasma membrane GFP

## References

Bao, S., Q. Wu, R.E. McLendon, Y. Hao, Q. Shi, A.B. Hjelmeland, M.W. Dewhirst, D.D. Bigner, and J.N. Rich. 2006. Glioma stem cells promote radioresistance by preferential activation of the DNA damage response. Nature 444:756–760.

Batlle, E., and H. Clevers. 2017. Cancer stem cells revisited. Nature medicine 23:1124–1134.

Bu, P., L. Wang, K.Y. Chen, T. Srinivasan, P.K. Murthy, K.L. Tung, A.K. Varanko, H.J. Chen, Y. Ai, S. King, S.M. Lipkin, and X. Shen. 2016. A miR-34a-Numb Feedforward Loop Triggered by Inflammation Regulates Asymmetric Stem Cell Division in Intestine and Colon Cancer. Cell stem cell 18:189–202.

Chen, G., J. Kong, C. Tucker-Burden, M. Anand, Y. Rong, F. Rahman, C.S. Moreno, E.G. Van Meir, C.G. Hadjipanayis, and D.J. Brat. 2014. Human Brat ortholog TRIM3 is a tumor suppressor that regulates asymmetric cell division in glioblastoma. Cancer research 74:4536–4548.

Chen, W., J. Dong, J. Haiech, M.C. Kilhoffer, and M. Zeniou. 2016. Cancer Stem Cell Quiescence and Plasticity as Major Challenges in Cancer Therapy. Stem cells international 2016:1740936.

Chumakova, A.P., M. Hitomi, E.P. Sulman, and J.D. Lathia. 2019. High-throughput automated single-cell imaging analysis reveals dynamics of glioblastoma stem cell population during state transition. Cytometry. Part A : the journal of the International Society for Analytical Cytology

Cicalese, A., G. Bonizzi, C.E. Pasi, M. Faretta, S. Ronzoni, B. Giulini, C. Brisken, S. Minucci, P.P. Di Fiore, and P.G. Pelicci. 2009. The tumor suppressor p53 regulates polarity of self-renewing divisions in mammary stem cells. Cell 138:1083–1095.

Clarke, J.L., A.M. Molinaro, J.J. Phillips, N.A. Butowski, S.M. Chang, A. Perry, J.F. Costello, A.A. DeSilva, J.E. Rabbitt, and M.D. Prados. 2014. A single-institution phase II trial of radiation, temozolomide, erlotinib, and bevacizumab for initial treatment of glioblastoma. Neuro-oncology 16:984–990.

Clarke, M.F., J.E. Dick, P.B. Dirks, C.J. Eaves, C.H. Jamieson, D.L. Jones, J. Visvader, I.L. Weissman, and G.M. Wahl. 2006. Cancer stem cells-perspectives on current status and future directions: AACR Workshop on cancer stem cells. Cancer research 66:9339–9344.

Cusulin, C., C. Chesnelong, P. Bose, M. Bilenky, K. Kopciuk, J.A. Chan, J.G. Cairncross, S.J. Jones, M.A. Marra, H.A. Luchman, and S. Weiss. 2015. Precursor States of Brain Tumor Initiating Cell Lines Are Predictive of Survival in Xenografts and Associated with Glioblastoma Subtypes. Stem cell reports 5:1–9.

Daynac, M., M. Chouchane, H.Y. Collins, N.E. Murphy, N. Andor, J. Niu, S.P.J. Fancy, W.B. Stallcup, and C.K. Petritsch. 2018. Lgl1 controls NG2 endocytic pathway to regulate oligodendrocyte differentiation and asymmetric cell division and gliomagenesis. Nature communications 9:2862.

Ganguly, D., M. Fan, C.H. Yang, B. Zbytek, D. Finkelstein, M.F. Roussel, and L.M. Pfeffer. 2018. The critical role that STAT3 plays in glioma-initiating cells: STAT3 addiction in glioma. Oncotarget 9:22095–22112.

Guichet, P.O., S. Guelfi, C. Ripoll, M. Teigell, J.C. Sabourin, L. Bauchet, V. Rigau, B. Rothhut, and J.P. Hugnot. 2016. Asymmetric Distribution of GFAP in Glioma Multipotent Cells. PloS one 11:e0151274.

Halatsch, M.E., U. Schmidt, J. Behnke-Mursch, A. Unterberg, and C.R. Wirtz. 2006. Epidermal growth factor receptor inhibition for the treatment of glioblastoma multiforme and other malignant brain tumours. Cancer treatment reviews 32:74–89.

Hu, Y., and G.K. Smyth. 2009. ELDA: extreme limiting dilution analysis for comparing depleted and enriched populations in stem cell and other assays. Journal of immunological methods 347:70–78.

Johnston, A.L., X. Lun, J.J. Rahn, A. Liacini, L. Wang, M.G. Hamilton, I.F. Parney, B.L. Hempstead, S.M. Robbins, P.A. Forsyth, and D.L. Senger. 2007. The p75 neurotrophin receptor is a central regulator of glioma invasion. PLoS biology 5:e212.

Knoblich, J.A. 2008. Mechanisms of asymmetric stem cell division. Cell 132:583–597.

Lathia, J.D., M. Hitomi, J. Gallagher, S.P. Gadani, J. Adkins, A. Vasanji, L. Liu, C.E. Eyler, J.M. Heddleston, Q. Wu, S. Minhas, A. Soeda, D.J. Hoeppner, R. Ravin, R.D. McKay, R.E. McLendon, D. Corbeil, A. Chenn, A.B. Hjelmeland, D.M. Park, and J.N. Rich. 2011. Distribution of CD133 reveals glioma stem cells self-renew through symmetric and asymmetric cell divisions. Cell death & disease 2:e200.

Liu, Y., L. Siles, X. Lu, K.C. Dean, M. Cuatrecasas, A. Postigo, and D.C. Dean. 2018. Mitotic polarization of transcription factors during asymmetric division establishes fate of forming cancer cells. Nature communications 9:2424.

Longo, F.M., and S.M. Massa. 2013. Small-molecule modulation of neurotrophin receptors: a strategy for the treatment of neurological disease. Nature reviews. Drug discovery 12:507–525.

Moitra, K., H. Lou, and M. Dean. 2011. Multidrug efflux pumps and cancer stem cells: insights into multidrug resistance and therapeutic development. Clinical pharmacology and therapeutics 89:491–502.

Mukherjee, S., C. Tucker-Burden, C. Zhang, K. Moberg, R. Read, C. Hadjipanayis, and D.J. Brat. 2016. Drosophila Brat and Human Ortholog TRIM3 Maintain Stem Cell Equilibrium and Suppress Brain Tumorigenesis by Attenuating Notch Nuclear Transport. Cancer research 76:2443–2452.

Obernier, K., A. Cebrian-Silla, M. Thomson, J.I. Parraguez, R. Anderson, C. Guinto, J. Rodas Rodriguez, J.M. Garcia-Verdugo, and A. Alvarez-Buylla. 2018. Adult Neurogenesis Is Sustained by Symmetric Self-Renewal and Differentiation. Cell stem cell 22:221–234 e228.

Osuka, S., and E.G. Van Meir. 2017. Overcoming therapeutic resistance in glioblastoma: the way forward. The Journal of clinical investigation 127:415–426.

Patel, A.P., I. Tirosh, J.J. Trombetta, A.K. Shalek, S.M. Gillespie, H. Wakimoto, D.P. Cahill, B.V. Nahed, W.T. Curry, R.L. Martuza, D.N. Louis, O. Rozenblatt-Rosen, M.L. Suva, A. Regev, and B.E. Bernstein. 2014. Single-cell RNA-seq highlights intratumoral heterogeneity in primary glioblastoma. Science 344:1396–1401.

Pyenta, P.S., D. Holowka, and B. Baird. 2001. Cross-correlation analysis of inner-leaflet-anchored green fluorescent protein co-redistributed with IgE receptors and outer leaflet lipid raft components. Biophysical journal 80:2120–2132.

Qazi, M.A., P. Vora, C. Venugopal, S.S. Sidhu, J. Moffat, C. Swanton, and S.K. Singh. 2017. Intratumoral heterogeneity: pathways to treatment resistance and relapse in human glioblastoma. Annals of oncology : official journal of the European Society for Medical Oncology 28:1448–1456.

Richardson, M.E., and D.W. Siemann. 1997. Tumor cell heterogeneity: impact on mechanisms of therapeutic drug resistance. International journal of radiation oncology, biology, physics 39:789–795.

Roper, K., D. Corbeil, and W.B. Huttner. 2000. Retention of prominin in microvilli reveals distinct cholesterol-based lipid micro-domains in the apical plasma membrane. Nature cell biology 2:582–592.

Sarkaria, J.N., B.L. Carlson, M.A. Schroeder, P. Grogan, P.D. Brown, C. Giannini, K.V. Ballman, G.J. Kitange, A. Guha, A. Pandita, and C.D. James. 2006. Use of an orthotopic xenograft model for assessing the effect of epidermal growth factor receptor amplification on glioblastoma radiation response. Clinical cancer research : an official journal of the American Association for Cancer Research 12:2264–2271.

Sartor, C.I., M.L. Dziubinski, C.L. Yu, R. Jove, and S.P. Ethier. 1997. Role of epidermal growth factor receptor and STAT-3 activation in autonomous proliferation of SUM-102PT human breast cancer cells. Cancer research 57:978–987.

Schonberg, D.L., T.E. Miller, Q. Wu, W.A. Flavahan, N.K. Das, J.S. Hale, C.G. Hubert, S.C. Mack, A.M. Jarrar, R.T. Karl, A.M. Rosager, A.M. Nixon, P.J. Tesar, P. Hamerlik, B.W. Kristensen, C. Horbinski, J.R. Connor, P.L. Fox, J.D. Lathia, and J.N. Rich. 2015. Preferential Iron Trafficking Characterizes Glioblastoma Stem-like Cells. Cancer cell 28:441–455.

Silver, D.J., and J.D. Lathia. 2018. Revealing the glioma cancer stem cell interactome, one niche at a time. The Journal of pathology 244:260–264.

Simons, K., and D. Toomre. 2000. Lipid rafts and signal transduction. Nature reviews. Molecular cell biology 1:31–39.

Srinivasan, T., J. Walters, P. Bu, E.B. Than, K.L. Tung, K.Y. Chen, N. Panarelli, J. Milsom, L. Augenlicht, S.M. Lipkin, and X. Shen. 2016. NOTCH Signaling Regulates Asymmetric Cell Fate of Fast-and Slow-Cycling Colon Cancer-Initiating Cells. Cancer research 76:3411–3421.

Sugiarto, S., A.I. Persson, E.G. Munoz, M. Waldhuber, C. Lamagna, N. Andor, P. Hanecker, J. Ayers-Ringler, J. Phillips, J. Siu, D.A. Lim, S. Vandenberg, W. Stallcup, M.S. Berger, G. Bergers, W.A. Weiss, and C. Petritsch. 2011. Asymmetry-defective oligodendrocyte progenitors are glioma precursors. Cancer cell 20:328–340.

Thakkar, J.P., T.A. Dolecek, C. Horbinski, Q.T. Ostrom, D.D. Lightner, J.S. Barnholtz-Sloan, and J.L. Villano. 2014. Epidemiologic and molecular prognostic review of glioblastoma. Cancer epidemiology, biomarkers & prevention : a publication of the American Association for Cancer Research, cosponsored by the American Society of Preventive Oncology 23:1985–1996.

Thorne, A.H., C. Zanca, and F. Furnari. 2016. Epidermal growth factor receptor targeting and challenges in glioblastoma. Neuro-oncology 18:914–918.

Tomasetti, C., and D. Levy. 2010. Role of symmetric and asymmetric division of stem cells in developing drug resistance. Proceedings of the National Academy of Sciences of the United States of America 107:16766–16771.

Tominaga, K., H. Minato, T. Murayama, A. Sasahara, T. Nishimura, E. Kiyokawa, H. Kanauchi, S. Shimizu, A. Sato, K. Nishioka, E.I. Tsuji, M. Yano, T. Ogawa, H. Ishii, M. Mori, K. Akashi, K. Okamoto, M. Tanabe, K.I. Tada, A. Tojo, and N. Gotoh. 2019. Semaphorin signaling via MICAL3 induces symmetric cell division to expand breast cancer stem-like cells. Proceedings of the National Academy of Sciences of the United States of America 116:625–630.

van den Bent, M.J., A.A. Brandes, R. Rampling, M.C. Kouwenhoven, J.M. Kros, A.F. Carpentier, P.M. Clement, M. Frenay, M. Campone, J.F. Baurain, J.P. Armand, M.J. Taphoorn, A. Tosoni, H. Kletzl, B. Klughammer, D. Lacombe, and T. Gorlia. 2009. Randomized phase II trial of erlotinib versus temozolomide or carmustine in recurrent glioblastoma: EORTC brain tumor group study 26034. Journal of clinical oncology : official journal of the American Society of Clinical Oncology 27:1268–1274.

Venkei, Z.G., and Y.M. Yamashita. 2018. Emerging mechanisms of asymmetric stem cell division. The Journal of cell biology 217:3785–3795.

Voelzke, W.R., W.J. Petty, and G.J. Lesser. 2008. Targeting the epidermal growth factor receptor in high-grade astrocytomas. Current treatment options in oncology 9:23–31.

Wang, L., P. Bu, Y. Ai, T. Srinivasan, H.J. Chen, K. Xiang, S.M. Lipkin, and X. Shen. 2016. A long non-coding RNA targets microRNA miR-34a to regulate colon cancer stem cell asymmetric division. eLife 5:

Wang, X., B.C. Prager, Q. Wu, L.J.Y. Kim, R.C. Gimple, Y. Shi, K. Yang, A.R. Morton, W. Zhou, Z. Zhu, E.A.A. Obara, T.E. Miller, A. Song, S. Lai, C.G. Hubert, X. Jin, Z. Huang, X. Fang, D. Dixit, W. Tao, K. Zhai, C. Chen, Z. Dong, G. Zhang, S.M. Dombrowski, P. Hamerlik, S.C. Mack, S. Bao, and J.N. Rich. 2018. Reciprocal Signaling between Glioblastoma Stem Cells and Differentiated Tumor Cells Promotes Malignant Progression. Cell stem cell 22:514–528 e515.

Xin, H.W., D.M. Hari, J.E. Mullinax, C.M. Ambe, T. Koizumi, S. Ray, A.J. Anderson, G.W. Wiegand, S.H. Garfield, S.S. Thorgeirsson, and I. Avital. 2012. Tumor-initiating label-retaining cancer cells in human gastrointestinal cancers undergo asymmetric cell division. Stem cells 30:591–598.

Zimdahl, B., T. Ito, A. Blevins, J. Bajaj, T. Konuma, J. Weeks, C.S. Koechlein, H.Y. Kwon, O. Arami, D. Rizzieri, H.E. Broome, C. Chuah, V.G. Oehler, R. Sasik, G. Hardiman, and T. Reya. 2014. Lis1 regulates asymmetric division in hematopoietic stem cells and in leukemia. Nature genetics 46:245–252.

