## Supplemental information for "Asymmetric Division Promotes Therapeutic Resistance in Glioblastoma Stem Cells"

#### SUPPLEMENTAL INFORMATION TITLES AND LEGENDS

**Supplemental Figure 1, related to Figure 1. A plasma membrane green fluorescence protein (PMGFP) reporter system allows the reliable evaluation of cell division mode and reveals functional differences in asymmetrically divided cells.**

**A** - Self-renewal capacity of T4121-PMGFP compared to non-transfected T4121 cells. N.S. indicates lack of significant difference.

**B** - Correlation between the percentage of PMGFP asymmetry at the time of mitosis and SOX2 percent asymmetry of daughter cells at the end of the time-lapse microscopy. Pearson's correlation coefficient was calculated and demonstrated a significant association between PMGFP reporter asymmetry and expression of the stem cell marker ($p = 0.03$).

**C** - Schematic work-flow of the FACS-based isolation of symmetrically and asymmetrically divided cells. **1**) T4121-PMGFP CSCs were synchronized in S-phase and labeled with CellTrace dye to monitor division. **2**) To increase the accuracy of the assay, cells were subjected to FACS sorting to obtain a population of cells with uniform PMGFP and CellTrace intensity. **3**) The cells were then released from thymidine into ACD-stimulating conditions (10% FBS) for 15 hours and then **4**) subjected to FACS: divided cells were separated by gating for cells with a CellTrace intensity that was half of the intensity of the undivided cells. Divided cells were then sorted for the top and bottom 5% of PMGFP intensity, which represents asymmetrically divided cells, as well as the PMGFP-mid population that represents symmetrically divided cells.

**Supplemental Figure 2, related to Figure 2. EGFR and p75NTR co-segregate during asymmetric cell division.**

**A** - Immunohistochemistry staining for EGFR in a human GBM specimen. A cell in metaphase exhibits asymmetric distribution of EGFR as defined by intensity threshold analysis (lower image).

**B** - Quantification of relative immunohistochemistry staining intensity on the dark side vs bright side of the metaphase cell membrane.

**C** - Immunofluorescence staining of a T4121 PMGFP intracranial tumor section showing an asymmetrically dividing cell as defined by a characteristic DNA condensation (blue) and alpha-tubulin (red)-positive mitotic spindle. Asymmetry of EGFR staining intensity corresponds to the asymmetry of PMGFP staining intensity.

**Supplemental Figure 3, related to Figure 4. Alteration of the p75NTR axis modifies response to EGFR inhibition.**

**A** - $IC_{50}$ of erlotinib (μM) in T4121-PMGFP cells expressing non-targeting (NT) or p75NTR knockdown shRNA (KD1, KD2, *** $p<0.000001$).

**B** - Survival analysis of animals with intracranial T4121-PMGFP tumors. NT indicates cells expressing non-targeting shRNA, and KD indicates cells expressing p75NTR knockdown shRNA. n = number of animals in each group; median survival and the p-value as determined by log-rank test comparing the vehicle and erlotinib groups are shown.

**C** - Survival analysis of an in vivo erlotinib dose escalation. Animals with intracranial T4121-PMGFP tumors were treated for 4 weeks with 0, 5, 15, 50 and 100 mg/kg of erlotinib. n = number of animals in each group; median survival and the p-value as determined by log-rank test comparing the vehicle and erlotinib groups are shown.

#### Supplemental figure 1 - related to Figure 1

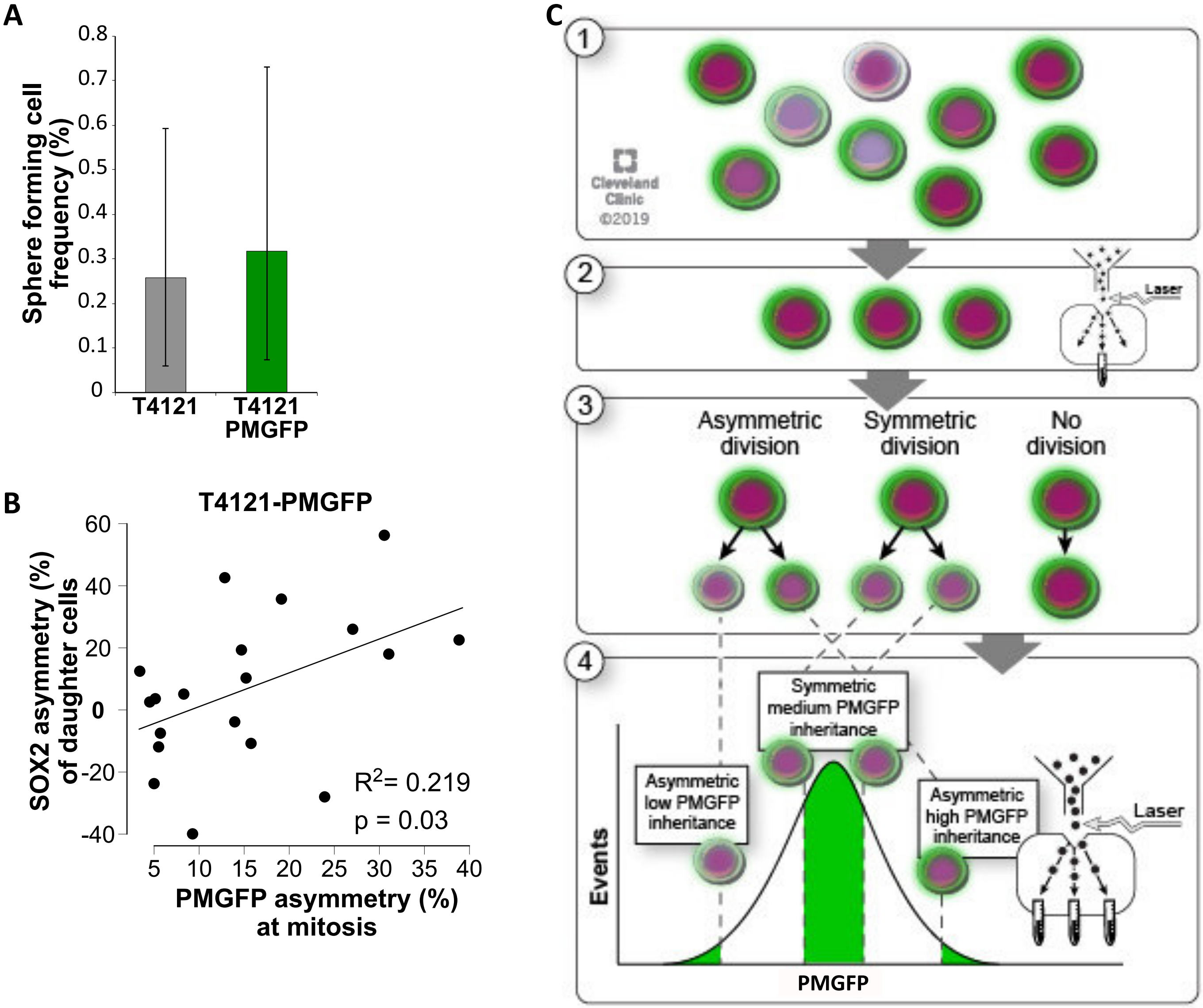

#### Supplemental figure 2 - related to Figure 2

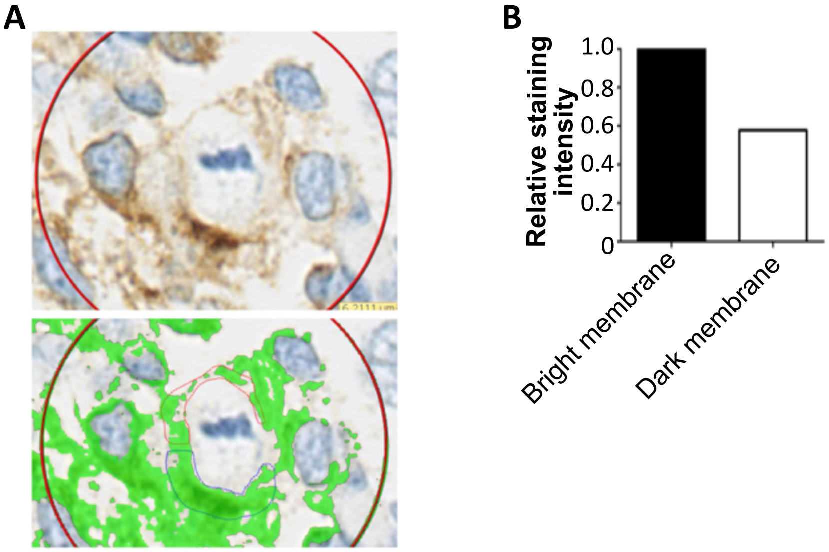


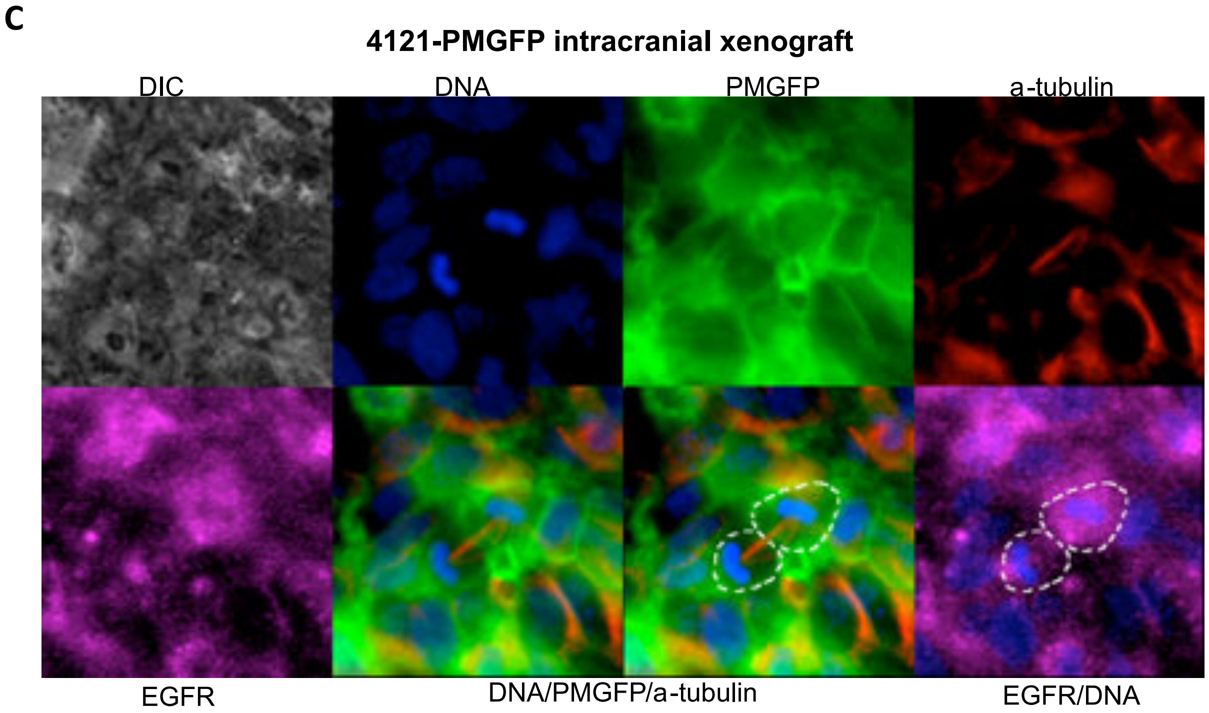

### Supplemental figure 3 - Figure 4

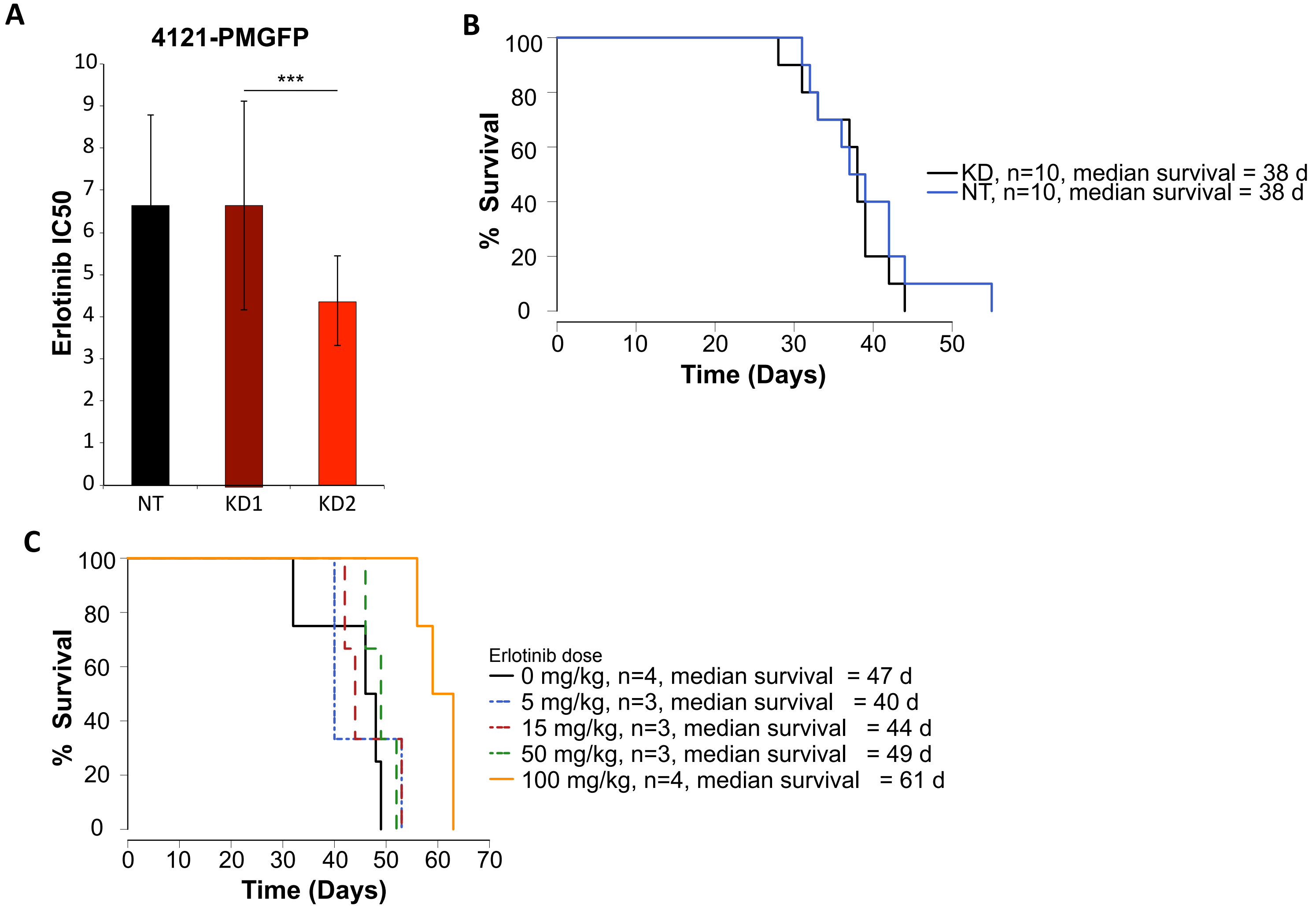
